# Causal contributions of the dorsolateral prefrontal cortex and temporoparietal junction to source and reality monitoring

**DOI:** 10.64898/2026.08.25.746632

**Authors:** C.M. Bates, L. Ring, M. Tolfrey, A.K. Martin

## Abstract

Source and reality monitoring enable individuals to distinguish the origins of remembered information, including whether information was self- or other-generated and whether it was perceived or imagined. Although the dorsolateral prefrontal cortex (dlPFC) and temporoparietal junction (TPJ) have been implicated in these processes, their independent causal contributions remain unclear. We investigated whether focal transcranial direct current stimulation (f-tDCS) of the left dlPFC and left TPJ differentially modulates source and reality monitoring. One hundred participants were randomly assigned to receive anodal or sham stimulation of the left dlPFC or TPJ before completing an episodic memory task manipulating agent (self, experimenter), context (spoken, imagined), and emotional valence (positive, negative). Discrimination sensitivity (*d*⍰) and response criterion (*c*) were examined separately. For source monitoring, stimulation interacted with context and cortical region: anodal dlPFC stimulation was associated with a greater spoken-imagined difference in self-experimenter discrimination than sham stimulation, whereas no equivalent context-dependent effect emerged following TPJ stimulation. For reality monitoring, stimulation effects also differed by cortical target, with reduced spoken-imagined discrimination following anodal relative to sham TPJ stimulation and no significant effect of dlPFC stimulation. These effects were not accompanied by corresponding stimulation effects on response criterion. Independent of stimulation, source discrimination was substantially greater for spoken than imagined information, while reality-monitoring sensitivity was enhanced for self-generated relative to experimenter-generated negative information. Together, these findings provide evidence that the dlPFC and TPJ make dissociable contributions to source and reality monitoring, while highlighting the importance of contextual and affective features in determining how the origins of memories are evaluated.

---

Source monitoring (SM) refers to the cognitive processes through which individuals infer the origins of remembered information (Johnson et al., 1993). One socially relevant dimension of source is agency: determining whether information originated from oneself or another person. Reality monitoring (RM) represents a specific form of source monitoring concerned with distinguishing internally generated experiences, such as thoughts or imagined events, from externally perceived experiences (Johnson & Raye, 1981). According to the Source Monitoring Framework, memories do not contain explicit labels specifying their origins; rather, source judgements are inferred from qualitative characteristics of the retrieved representation, including perceptual and contextual information, semantic and affective detail, and records of the cognitive operations engaged when the memory was formed (Johnson et al., 1993).

Together, SM and RM allow individuals to reconstruct different aspects of the origin of remembered information, including whether it was associated with oneself or another person and whether it was externally perceived or internally generated, supporting coherent representations of past experience, the self, and the external world (Gallagher, 2000; Johnson et al., 1993; Johnson & Raye, 1981). Failures in source and reality monitoring (SRM) are associated with hallucinatory symptoms across the psychosis spectrum (Damiani et al., 2022), dissociation (Suter et al., 2026), and Alzheimer’s disease (El Haj et al., 2012), highlighting the importance of understanding their underlying cognitive and neural mechanisms.

Within episodic memory, source and reality judgements require the reconstruction and evaluation of features encoded as part of a past event, including perceptual, contextual, cognitive, and agent-related information. This can be examined using episodic memory paradigms in which different attributes of an event are manipulated at encoding and subsequently retrieved. For example, Kwon et al. (2022) asked participants to encode objects associated with either self-versus experimenter-generated actions (SM) or imagined versus perceived movements (RM), before later identifying the original source and reality status of each memory. Participants exhibited an externalisation bias across both SM and RM, with poorer accuracy when attributing internally generated than externally generated events.

Relatedly, previous word-based episodic memory studies have demonstrated enhanced source monitoring for self-related information when words are encoded in relation to the self rather than another person (Bates, Dias Salgado, et al., 2026; Bates, Lingawi, et al., 2026; Chirtop et al., 2025; Durbin et al., 2017). Differences across these paradigms may reflect how self-related processing is operationalised. Self-referential encoding can promote elaborative processing and strengthen the binding of information to its source, whereas discriminating physically enacted self- from other-generated events requires recovery of agency-specific episodic features and may therefore be more susceptible to externalisation errors. Thus, the ability to identify the source of an episodic memory is likely to depend on the diagnostic features available when its origin is reconstructed.

Emotional content may further influence the diagnostic information available for source attribution. Emotionally salient information typically receives enhanced attentional allocation and contextual binding during encoding (e.g., Dent & Martin, 2023; Leshikar & Duarte, 2012; Narhi-Martinez et al., 2023). Consistent with this, recent evidence suggests that self- and close-other advantages in source memory are particularly evident for positive information (Bates & Martin, 2026). Emotional content can also influence the likelihood of confusing imagined and perceived events. Negative and highly arousing information can enhance the encoding or retention of perceptual and contextual details, potentially providing more diagnostic information during subsequent reality monitoring. For example, imagined negative events may be less likely to be misattributed as externally perceived than neutral events, suggesting that emotional arousal can strengthen memory for source-specific features (Kensinger et al., 2007). Source and reality judgements may therefore depend not only on whether information was self- or other-generated or perceived or imagined, but also on the affective characteristics of the remembered event.

If source and reality judgements depend on retrieving and evaluating different features of an episodic representation, partially dissociable neural systems may support these monitoring processes. Two particularly strong candidates are the temporoparietal junction (TPJ) and dorsolateral prefrontal cortex (dlPFC), which may contribute differently to reconstructing the origin of episodic memories. The TPJ has been implicated in distinguishing internally generated from externally derived information and in representing self–other distinctions (Eddy, 2016; Pryke et al., 2025; Quesque & Brass, 2019; Sperduti et al., 2011; Sun et al., 2023; Wen et al., 2021). The TPJ is also associated with perspective-taking, empathising, and distinguishing and switching between one’s own mental and affective states from those of others (Eliášová et al., 2026; Martin et al., 2019, 2020; Santiesteban et al., 2012; Sowden & Catmur, 2015). Much of this evidence, particularly that concerning self–other control and perspective-taking, has implicated the right TPJ. However, TPJ contributions appear to vary according to the cognitive process under investigation, and the left TPJ may be particularly relevant when source judgements involve verbal information (Perret et al., 2024). More broadly, neuroimaging studies consistently report TPJ engagement during successful recollection, particularly when retrieved memories contain vivid contextual or self-relevant information (Cabeza et al., 2008; Vilberg & Rugg, 2008). Because SRM requires the retrieval and evaluation of contextual features that distinguish self from other and internally generated from externally perceived events, the TPJ may contribute to representing the mnemonic information on which these judgements are based. Critically, direct causal evidence supports a role for the left TPJ in verbal reality monitoring: Mondino et al. (2016) demonstrated that anodal stimulation of the left TPJ increased externalisation bias, such that participants were more likely to misattribute self-generated events as externally perceived, while leaving SM unaffected. Thus, although broader self–other processing has often been associated with the right TPJ, evidence from verbal source-monitoring paradigms provides a specific rationale for targeting the left TPJ in the present study.

The dlPFC may make a complementary contribution through the strategic retrieval and evaluation of source information. The lateral prefrontal cortex has been implicated in higher-order monitoring processes involved in strategic retrieval, executive control, and post-retrieval evaluation (Fletcher et al., 1998; Friedman & Robbins, 2022). Neuroimaging evidence indicates that lateral prefrontal regions are recruited when retrieval requires the recollection and evaluation of source information rather than simpler item recognition, consistent with a role in monitoring retrieved information in relation to current task (Dobbins et al., 2004; Rugg et al., 2003; Turner et al., 2008). A recent systematic review of non-invasive brain-stimulation studies provides complementary causal evidence, identifying lateral prefrontal regions, including the left rostrolateral PFC and dlPFC, together with bilateral temporoparietal cortices, as contributors to internal source monitoring (Perret et al., 2024). Reality-monitoring errors have also been associated with reduced prefrontal activation (Simons et al., 2017). Increased functional connectivity between frontal and temporal regions has been demonstrated when comparing self- with other-related source monitoring (Wang et al., 2011), suggesting that successful source attribution may depend on interactions between regions representing mnemonic information and frontal systems involved in evaluating that information. Such evaluative processes may be especially important when retrieved events contain multiple diagnostic features that can be used to infer their source. For example, overtly spoken events contain auditory and voice-identity information and, when self-produced, additional articulatory, motor, and proprioceptive features. Strategic evaluation of these features may facilitate discrimination between self- and other-generated memories or between spoken and imagined events, providing a potential mechanism through which the dlPFC contributes to SRM.

Brain-stimulation studies provide further causal support for prefrontal and temporoparietal involvement in source monitoring. In their systematic review of 23 non-invasive brain-stimulation studies, Perret et al. (2024) identified a lateral prefrontal– temporoparietal circuit involved in internal source monitoring and a medial prefrontal– temporoparietal circuit involved in reality monitoring. Importantly, the authors also emphasised that these subprocesses are unlikely to rely on entirely distinct neural systems, given their shared dependence on self-generation, imagery, perceptual information, and speech-related processes. Individual stimulation studies similarly suggest that modulating frontotemporal systems can alter source-monitoring performance. Kusztrits et al. (2021) combined cathodal stimulation of the prefrontal cortex with anodal stimulation of the superior temporal gyrus, resulting in improved SM while leaving RM unaffected. More recently, Sharma et al. (2024) used conventional bipolar tDCS, applying anodal stimulation over the left superior temporal gyrus and cathodal stimulation over the left dlPFC before participants completed reality monitoring (Hear–Imagine), internal source monitoring (Say–Imagine), and external source monitoring (Real–Virtual) tasks. Active stimulation improved internal source monitoring and, to a lesser extent, reality monitoring, whereas external source monitoring remained unaffected. However, because these studies simultaneously stimulated multiple cortical regions, the specific contribution of prefrontal relative to temporal regions cannot be determined. Moreover, conventional bipolar tDCS produces relatively diffuse current flow between relatively large electrodes. Sharma et al. therefore noted that the strongest electric field may have been centred over intervening regions, including Broca’s area, rather than being confined to the intended superior temporal and prefrontal targets.

Although converging neuroimaging and brain-stimulation evidence therefore implicates lateral prefrontal and temporoparietal regions in SRM (Perret et al., 2024), their independent causal contributions remain unclear. Previous stimulation studies relevant to SRM have either targeted a single cortical region (Mondino et al., 2016) or simultaneously modulated frontal and temporal regions (Kusztrits et al., 2021; Sharma et al., 2024), limiting the extent to which behavioural effects can be attributed to a specific neural target. Furthermore, these previous studies employed conventional tDCS, for which relatively diffuse current flow complicates anatomical interpretation. The present study addressed these limitations using focal transcranial direct current stimulation (f-tDCS) to independently manipulate the left dlPFC and left TPJ. Previous computational current modelling using identical electrode parameters confirmed that stimulation was concentrated over the intended cortical targets (see Pryke et al., 2025).

Participants received either anodal or sham f-tDCS over the left dlPFC or left TPJ before completing an episodic SRM task that independently manipulated agency (self vs. experimenter), reality (spoken vs. imagined), and emotional valence (positive vs. negative). By comparing the two cortical targets using the same task, stimulation protocol, and outcome measures, this design provided a direct test of whether the dlPFC and TPJ make dissociable causal contributions to retrieving and evaluating different features of an episodic event. Based on previous evidence, we hypothesised that stimulation effects would differ according to cortical target, with TPJ stimulation expected to exert a greater influence on disrupting RM. We also hypothesized that dlPFC stimulation would facilitate source monitoring. Given evidence that emotional content influences source attribution, we additionally examined whether source and reality monitoring differed according to the valence of the remembered information.

## Methods

### Participants

An a priori power analysis indicated that a minimum sample of 96 participants was required to detect a medium-sized effect (Cohen’s *f* = 0.25) with 80% power at α = .05. A total of 100 undergraduate students from the University of Kent were recruited through the university’s research participation scheme in exchange for course credit. Participants were randomly assigned to one of four groups in a 2 (Region: dlPFC, TPJ) × 2 (Stimulation: Anodal, Sham) between-subjects design, resulting in 25 participants per condition. All participants had normal or corrected-to-normal vision, no psychiatric diagnoses or neurological conditions, and no electrical medical equipment on their person that could not be removed. Ethical approval was granted by the Psychology Ethics Committee at the University of Kent.

### Design and analysis

The current study employed a series of mixed-design ANOVAs, with comparable analyses conducted separately for Reality Monitoring and Source Monitoring. For each dependent variable, discrimination sensitivity (*d*⍰) and response criterion (*c*) were analysed using the same factorial design.

For Reality Monitoring, a 2 (Stimulation: sham, anodal) × 2 (Region: PFC, TPJ) × 2 (Agent: self, experimenter) × 2 (Valence: positive, negative) mixed ANOVA was conducted on *d*⍰ scores. Stimulation and Region were entered as between-participants factors, whereas Agent and Valence were entered as within-participants factors. An identical mixed ANOVA was conducted on response criterion (*c*).

For Source Monitoring, a 2 (Stimulation: sham, anodal) × 2 (Region: PFC, TPJ) × 2 (Context: spoken, imagined) × 2 (Valence: positive, negative) mixed ANOVA was conducted on *d*⍰ scores. Stimulation and Region were entered as between-participants factors, whereas Context and Valence were entered as within-participants factors. An identical mixed ANOVA was conducted on response criterion (*c*).

Generalised eta-squared (*η^2^G*) was used as the measure of effect size for omnibus ANOVA effects. For significant interactions, follow-up comparisons were conducted to characterise the pattern of effects. Paired-samples *t*-tests were used for within-participant comparisons, and independent-samples *t*-tests were used for comparisons between the anodal and sham stimulation groups. Bonferroni corrections were applied separately within each family of two follow-up comparisons within each brain region, with both uncorrected and corrected *p*-values reported. Cohen’s *d* was reported for follow-up comparisons.

Additional analyses were conducted to assess potential group differences in mood, adverse effects, and age. Changes in positive and negative mood, as measured using the Visual Analogue Mood Scale (VAMS), were analysed using ANOVA. Adverse effects and participant age were also assessed using ANOVA to examine whether these measures differed as a function of the experimental group factors. The effectiveness of participant blinding was assessed separately within each brain region using exact binomial tests, comparing the proportion of participants who correctly identified their stimulation condition with chance performance (50%).

The presentation of all within-participants factors was randomised across trials.

### Signal-detection measures

The preregistered primary outcomes were discrimination sensitivity (dN) and response criterion (c). For each analysis, one source category was designated as the signal and the alternative source category formed the noise distribution.

For source monitoring, dN quantified discrimination between self- and experimenter-generated items separately within each Context × Valence condition. Self-generated items were designated as the signal. Hits were therefore defined as self responses to self-generated items, whereas false alarms were defined as self responses to experimenter-generated items. Experimenter responses to self-generated items constituted misses, and experimenter responses to experimenter-generated items constituted correct rejections.

For reality monitoring, dN quantified discrimination between spoken and imagined items separately within each Agent × Valence condition. Spoken items were designated as the signal. Hits were therefore defined as spoken responses to spoken items, whereas false alarms were defined as spoken responses to imagined items. Imagined responses to spoken items constituted misses, and imagined responses to imagined items constituted correct rejections. See Table 1. Details pertaining to the sensitivity analyses for this specific task are provided in detail elsewhere (Bates, Lingawi, et al., 2026).

**Table 1.** Classification of responses used to calculate signal-detection measures.

|  | True item category | Participant response | Classification |
| --- | --- | --- | --- |
| **Source Monitoring** | Self | Self | Hit |
|  | Self | Experimenter | Miss |
|  | Experimenter | Self | False alarm |
|  | Experimenter | Experimenter | Correct rejection |
| **Reality Monitoring** | Spoken | Spoken | Hit |
|  | Spoken | Imagined | Miss |
|  | Imagined | Spoken | False alarm |
|  | Imagined | Imagined | Correct rejection |

## Materials

### Hardware / Software

The cognitive task was presented through PsychoPy3^©^ (Peirce et al., 2019) and completed using a standard computer monitor, mouse, and keyboard. All questionnaires and demographic information were provided using Qualtrics (2019).

### Measures

All participants completed a tDCS safety screening questionnaire to ensure eligibility to participate. The Visual Analogue of Mood Scale (VAMS; Folstein & Luria, 1973) was employed to assess changes in both positive and negative moods following stimulation (anodal and sham). Positive mood was calculated by combining ratings of happiness and energy, whereas negative mood was calculated by combining ratings of anger, tiredness, tension, confusion, and sadness. For each mood dimension, a change score was calculated by subtracting the pre-stimulation score from the post-stimulation score. Adverse effects of stimulation were measured for both sham and anodal groups using an adapted questionnaire from Brunoni et al. (2011).

### Source-Reality Monitoring in Episodic Memory Task

In the current study, words were encoded under four conditions: spoken aloud by the experimenter, spoken aloud by the participant, imagined as spoken by the participant, or imagined as spoken by the experimenter. The reality component indexed whether the event was externally generated (overt speech) or internally generated (imagined speech), whereas the source component indexed the agent to whom the word was attributed at encoding (self vs. experimenter), irrespective of reality status. After encoding all 80 words, participants completed an unrelated ∼20-minute filler task, followed by a memory test requiring cued-recall of both the encoding context (real vs. imagined) and the agent (self vs. experimenter). Stimuli were selected from Warriner et al. (2013) and categorised as positive or negative based on whether the valence score was under or over 5 and categories were matched on arousal. Please refer to Figure 1 for a visual representation of the experimental trials.

**Figure 1.**
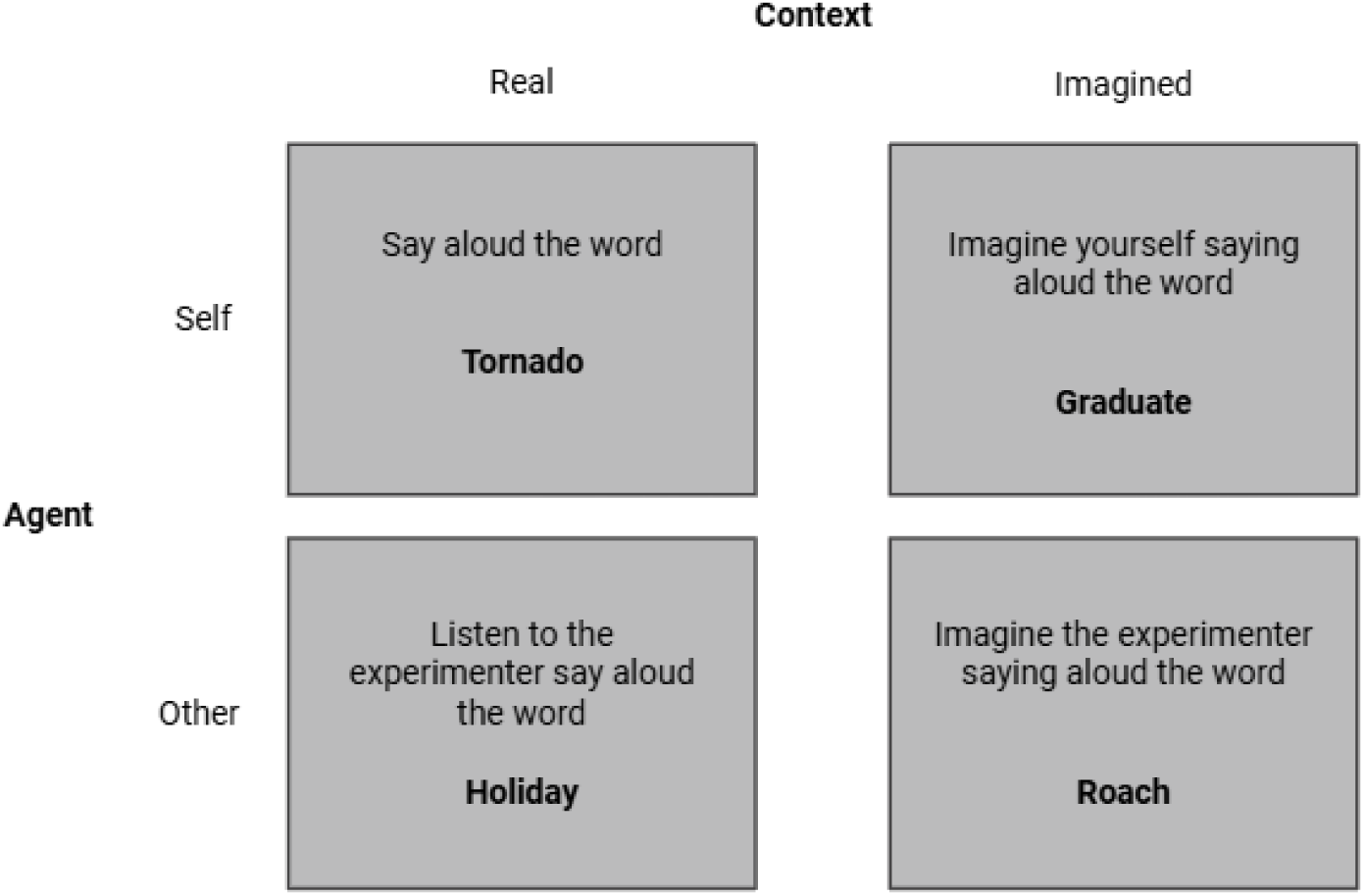
Visual representation of the experimental trails.

### Focal tDCS

The f-tDCS setup included a single-channel direct current stimulator (DC-Stimulator Plus, NeuroConn), with a central electrode (2.5cm diameter) and return electrode (inner/outer diameter of 7.5cm and 9.8cm, respectively) attached using adhesive electroconductive gel and an EEG cap. The central electrode determines stimulation polarity, in this case anodal, and the return electrode constrains electrical current to the target region, preserving spatial focality (Gbadeyan et al., 2016; Niemann et al., 2024). During anodal sessions, transcranial stimulation was delivered at 1 mA for 20 minutes, with a 5-second ramp-up and ramp-down at the beginning and end of stimulation. In the sham condition, stimulation was applied for 40 seconds, including the same ramp-up and ramp-down procedure, to mimic the initial sensation of stimulation while avoiding any lasting neurophysiological effects. Stimulation condition (anodal or sham) was randomised across participants. The study employed a double-blind design, whereby neither the participant nor the researcher administering the stimulation was aware of the stimulation condition. Blinding was achieved using the study mode function of the NeuroConn stimulator, which required the entry of anonymised codes corresponding to pre-programmed stimulation protocols. The l-dlPFC was located by finding 15% of the distance from the Fz towards the Fpz, whilst the l-TPJ was estimated at CP5. Previous current-flow modelling using identical electrode configurations demonstrated focal current delivery to the intended cortical targets (Pryke et al., 2025), please see Figure 2.

**Figure 2.**
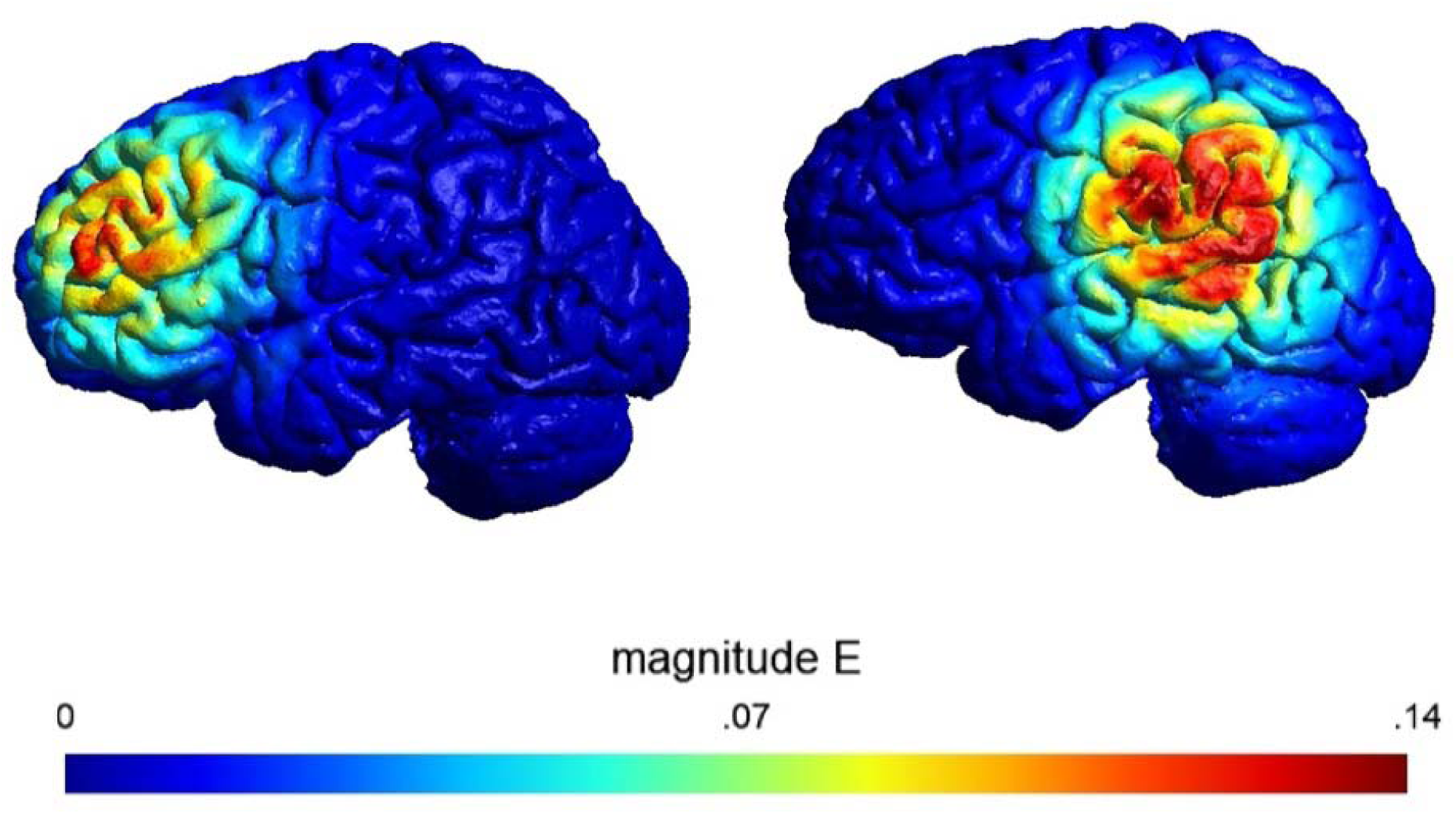
Illustrative current-flow model for the stimulation montage used in the present study.

## Procedure

Participants first completed a safety screening to confirm eligibility and absence of contraindications for f-tDCS. Eligible participants were then directed to Qualtrics to access an information sheet and provided informed written consent. They then completed a demographic questionnaire (age, gender) and the VAMS. Electrodes were subsequently positioned according to condition. Participants were informed about potential sensations and then completed the cognitive task described above. Following the task, participants completed the VAMS, an adverse effects measure, and were asked to guess whether they received active or sham stimulation. Participants were then debriefed and reminded of their right to withdraw their data.

## Results

### Demographics

The final sample consisted of 100 participants, with 25 participants allocated to each of the four experimental conditions: anodal stimulation of the prefrontal cortex (PFC), sham stimulation of the PFC, anodal stimulation of the temporoparietal junction (TPJ), and sham stimulation of the TPJ. Both the PFC and TPJ cohorts consisted of 20 women and 5 men in each stimulation condition.

Participants had a mean age of 19.48 years (SD = 1.71) in the PFC anodal condition, 19.72 years (SD = 2.81) in the PFC sham condition, 19.12 years (SD = 1.83) in the TPJ anodal condition, and 19.68 years (SD = 1.86) in the TPJ sham condition. A 2 × 2 ANOVA indicated that age did not significantly differ according to stimulation condition, *F*(1, 96) = 0.91, *p* = .343, η^2^p = .009, or brain region, *F*(1, 96) = 0.23, *p* = .635, η^2^p = .002. There was also no significant interaction between stimulation condition and brain region, *F*(1, 96) = 0.15, *p* = .704, η^2^p = .002.

### Source Monitoring

Means and standard deviations across all conditions are provided in Table 2.

**Table 2.** Means and Standard Deviations for *d*⍰ and Criterion (*c*) Across Brain Region and Stimulation Conditions.

| *Task / condition* | *PFC Sham* | *PFC Anodal* | *TPJ Sham* | *TPJ Anodal* |
| --- | --- | --- | --- | --- |
|  | *M (SD)* | *M (SD)* | *M (SD)* | *M (SD)* |
| **Source monitoring: $d'$** |  |  |  |  |
| *Imagined – Neg* | *0.55 (0.67)* | *0.49 (0.89)* | *0.63 (0.66)* | *0.47 (0.69)* |
| *Imagined – Pos* | *0.66 (0.53)* | *0.64 (0.81)* | *0.68 (0.53)* | *0.48 (0.58)* |
| *Spoken– Neg* | *1.49 (0.95)* | *1.85 (0.91)* | *1.56 (0.90)* | *1.35 (0.67)* |
| *Spoken– Pos* | *1.23 (0.81)* | *1.73 (0.98)* | *1.73 (0.88)* | *1.23 (0.70)* |
| **Source monitoring: c** |  |  |  |  |
| *Imagined – Neg* | *0.21 (0.36)* | *0.28 (0.34)* | *0.23 (0.46)* | *0.26 (0.45)* |
| *Imagined – Pos* | *0.11 (0.35)* | *0.17 (0.37)* | *0.18 (0.28)* | *0.17 (0.42)* |
| *Spoken– Neg* | *–0.01 (0.31)* | *0.03 (0.34)* | *0.13 (0.26)* | *0.12 (0.33)* |
| *Spoken– Pos* | *0.39 (0.28)* | *0.24 (0.32)* | *0.31 (0.37)* | *0.19 (0.23)* |
| **Reality monitoring: $d'$** |  |  |  |  |
| *Self – Neg* | *1.49 (0.75)* | *1.65 (0.84)* | *1.75 (0.63)* | *1.39 (0.58)* |
| *Self – Pos* | *1.32 (0.81)* | *1.25 (0.85)* | *1.61 (0.89)* | *1.18 (0.80)* |
| *Experimenter – Neg* | *0.71 (0.57)* | *0.94 (0.97)* | *1.03 (0.68)* | *0.92 (0.74)* |
| *Experimenter – Pos* | *1.07 (0.71)* | *1.27 (0.73)* | *1.52 (0.73)* | *1.08 (0.67)* |
| **Reality monitoring: c** |  |  |  |  |
| *Self – Neg* | *0.29 (0.33)* | *0.20 (0.40)* | *0.27 (0.37)* | *0.27 (0.35)* |
| *Self – Pos* | *0.49 (0.33)* | *0.39 (0.40)* | *0.38 (0.34)* | *0.40 (0.39)* |
| *Experimenter – Neg* | *0.55 (0.35)* | *0.26 (0.46)* | *0.36 (0.31)* | *0.38 (0.37)* |
| *Experimenter – Pos* | *0.42 (0.36)* | *0.28 (0.50)* | *0.30 (0.40)* | *0.39 (0.23)* |

### Discrimination sensitivity (*d*⍰)

The Context × Region × Stimulation interaction was significant, *F*(1, 96) = 6.03, *p* = .016, *η^2^G* = .011. To characterise this interaction, the Context × Stimulation interaction was examined separately within each region. Within the PFC condition, the Context × Stimulation interaction was significant, *F*(1, 48) = 5.37, *p* = .025, *η^2^G* = .028.

To decompose the PFC Context × Stimulation interaction, independent-samples *t*-tests compared the anodal and sham groups separately for spoken and imagined items. The effect of Stimulation was not significant for either spoken items, *t*(47.98) = 1.89, *p* = .065, *p*_bonf_ = .130, *d* = 0.53, or imagined items, *t*(44.16) = −0.24, *p* = .810, *p*_bonf_ = 1.00, *d* = −0.07. Thus, the significant interaction reflected a greater spoken–imagined difference in source-discrimination sensitivity following anodal than sham stimulation of the PFC (*M*diff Anodal = 1.23, *M*diff Sham = 0.75), primarily due to improved source discrimination for spoken words. The Context × Stimulation interaction was not significant within the TPJ condition, *F*(1, 48) = 1.09, *p* = .302, *p*_bonf_ = .605, *η^2^G* = .006. See Figure 3.

**Figure 3.**
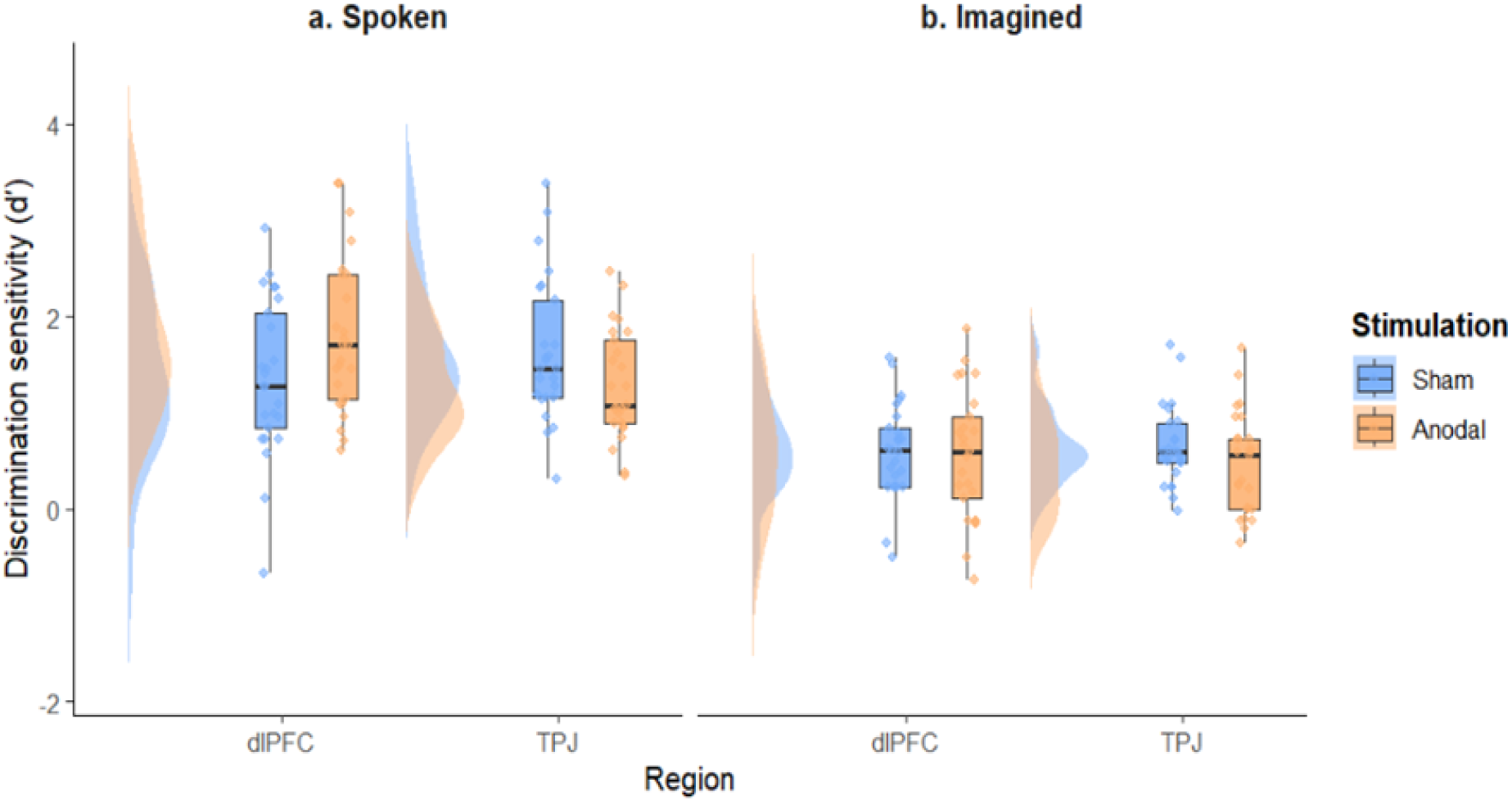
Stimulation effects on source monitoring sensitivity depending on brain region and spoken or imagined context.

There was also a significant main effect of Context, *F*(1, 96) = 205.80, *p* < .001, *η^2^G* = .280, with greater discrimination between spoken and imagined items.

There were no significant main effects of Valence, *F*(1, 96) < 0.01, *p* = .968, *η^2^G* < .001, Region, *F*(1, 96) = 0.32, *p* = .574, *η^2^G* = .002, or Stimulation, *F*(1, 96) = 0.11, *p* = .743, *η^2^G* < .001.

All remaining interactions were nonsignificant (see Supplementary Table 1).

### Response criterion (*c*)

There was no Context × Region × Stimulation interaction for *c*, *F*(1, 96) = 0.15, *p* = .695, *η^2^G* < .001. Thus, there was no evidence that the corresponding stimulation effect on *d*⍰ was attributable to a differential shift in response criterion.

There was a significant main effect of Valence, *F*(1, 96) = 3.96, *p* = .049, *η^2^G* = .008, with a more conservative self-response criterion for positive (*M* = 0.22, *SD* = 0.25) than negative words (*M* = 0.16, *SD* = 0.28). This effect was qualified by a significant Context × Valence interaction, *F*(1, 96) = 28.10, *p* < .001, *η^2^G* = .047.

Bonferroni-corrected paired comparisons indicated that, for negative words, *c* was significantly lower for spoken (*M* = 0.07, *SD* = 0.31) than imagined items (*M* = 0.25, *SD* = 0.40), *t*(99) = −4.01, *p* < .001, *p*_bonf_ < .001, *d* = −0.40. Conversely, for positive words, *c* was significantly higher for spoken (*M* = 0.28, *SD* = 0.31) than imagined items (*M* = 0.16, *SD* = 0.35), *t*(99) = 2.83, *p* = .006, *p*_bonf_ = .011, *d* = 0.28. The interaction therefore reflected a crossover in response criterion: participants showed a more liberal self-response criterion for spoken than imagined negative words, but a more conservative self-response criterion for spoken than imagined positive words.

All remaining main effects and interactions were nonsignificant (see Supplementary Table 2).

### Reality Monitoring

Means and standard deviations across all conditions are provided in Table 2.

### Discrimination sensitivity (*d*⍰)

There was a significant Region × Stimulation interaction, *F*(1, 96) = 4.20, *p* = .043, *η^2^G* = .025. Stimulation to the TPJ resulted in reduced discrimination sensitivity, *t*(47.76) = −2.19, *p* = .034, *d* = −0.62, whereas stimulation to the PFC had no significant effect, *t*(46.92) = 0.78, *p* = .439, *d* = 0.22 (see Figure 4.).

**Figure 4.**
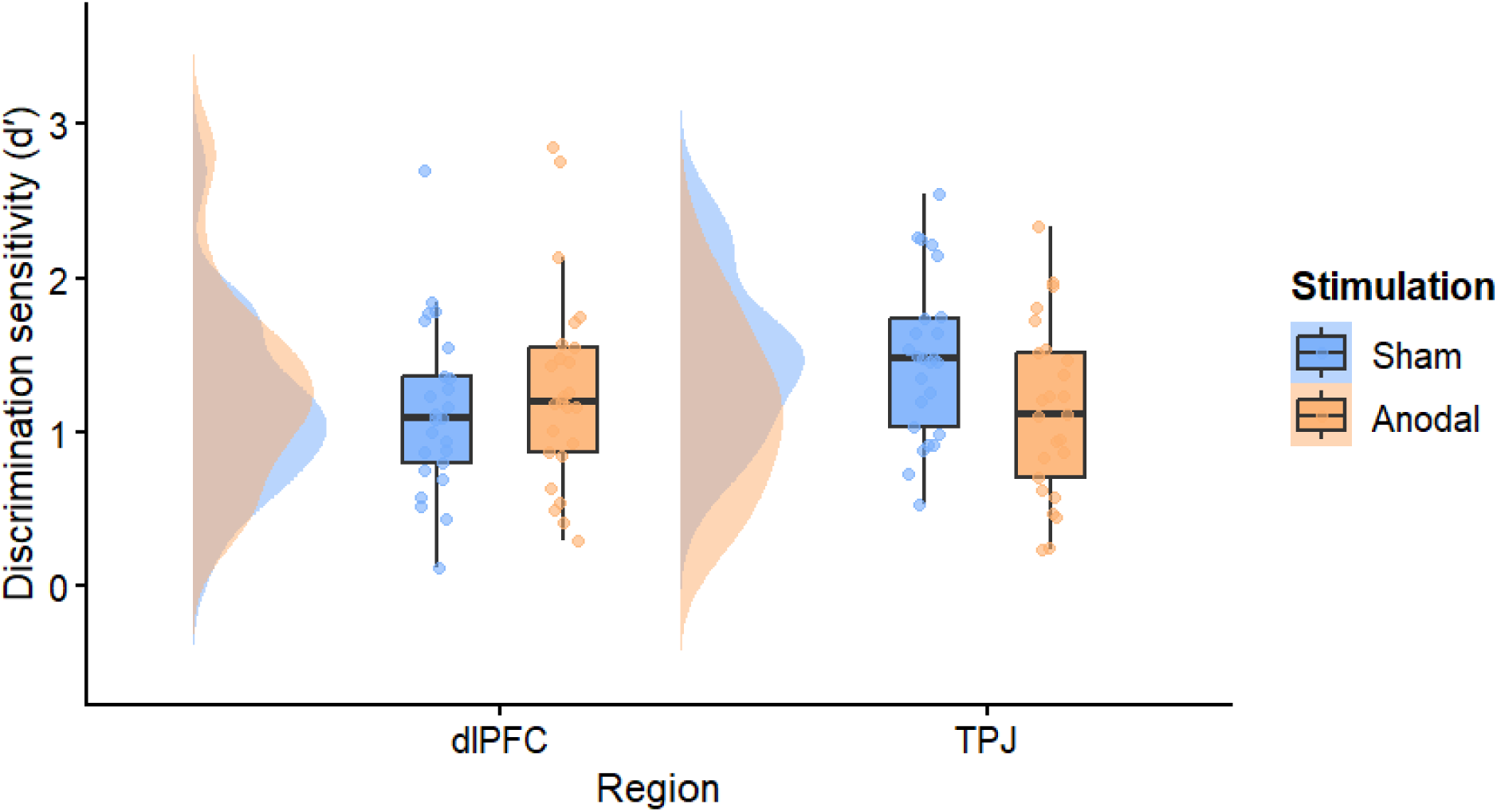
Stimulation effects on reality monitoring sensitivity by brain region.

There was also a significant main effect of Agent, *F*(1, 96) = 41.61, *p* < .001, *η^2^G* = .065, with greater discrimination sensitivity for self-generated (*M* = 1.45, *SD* = 0.67) than experimenter-generated words (*M* = 1.07, *SD* = 0.64). This effect was qualified by a significant Agent × Valence interaction, *F*(1, 96) = 29.11, *p* < .001, *η^2^G* = .035.

Bonferroni-corrected paired comparisons indicated that, for negative words, discrimination sensitivity was significantly greater for self-generated (*M* = 1.57, *SD* = 0.71) than experimenter-generated items (*M* = 0.90, *SD* = 0.75), *t*(99) = 8.35, *p* < .001, *p*_bonf_ < .001, *d* = 0.84. In contrast, sensitivity for positive words did not differ between self-generated (*M* = 1.34, *SD* = 0.84) and experimenter-generated items (*M* = 1.23, *SD* = 0.72), *t*(99) = 1.34, *p* = .182, *p*_bonf_ = .365, *d* = 0.13. Thus, the Agent × Valence interaction reflected greater reality-monitoring sensitivity for self- than experimenter-generated information specifically for negative words.

No other main effects or interactions were significant (see Supplementary Table 3).

### Response criterion (*c*)

No significant Region × Stimulation interaction was observed for response criterion, *F*(1, 96) = 3.09, *p* = .082, *η^2^G* = .017, indicating no evidence that stimulation differentially affected response criterion across the two regions.

There was, however, a significant Agent × Valence interaction, *F*(1, 96) = 14.87, *p* < .001, *η^2^G* = .018. Bonferroni-corrected paired comparisons indicated that, for negative words, *c* was significantly lower for self-generated (*M* = 0.26, *SD* = 0.36) than experimenter-generated items (*M* = 0.39, *SD* = 0.38), *t*(99) = −3.32, *p* = .001, *p*_bonf_ = .003, *d* = −0.33. This indicated a more liberal tendency to classify negative items as spoken when they had been self-generated. For positive words, *c* did not differ significantly between self-generated (*M* = 0.41, *SD* = 0.36) and experimenter-generated items (*M* = 0.35, SD = 0.38), *t*(99) = 1.76, *p* = .082, *p*_bonf_ = .163, *d* = 0.18.

Thus, the self-generated advantage in reality-monitoring sensitivity for negative words was accompanied by a more liberal spoken-response criterion for those items.

No other main effects or interactions were significant (see Supplementary Table 4).

### Stimulation Blinding

Blinding was assessed by comparing participants’ guesses of their stimulation condition with their actual condition separately for each brain region. In the PFC condition, 23 of 50 participants (46%) correctly identified their stimulation condition, which did not differ significantly from chance (50%), exact binomial p = .672. Similarly, in the TPJ condition, 28 of 50 participants (56%) correctly identified their stimulation condition, which also did not differ significantly from chance, exact binomial *p* = .480. These findings indicate that participants were unable to identify their stimulation condition above chance levels in either brain region, supporting successful blinding.

### Mood Change

Participants’ change in negative mood, as measured by the VAMS, was also examined across conditions. Mean negative mood change was −32.44 (SD = 76.89) in the PFC anodal condition, −16.88 (SD = 78.56) in the PFC sham condition, −31.44 (SD = 45.16) in the TPJ anodal condition, and −7.08 (SD = 72.94) in the TPJ sham condition. A 2 × 2 ANOVA found no significant main effect of stimulation condition, *F*(1, 96) = 2.05, *p* = .156, η²p = .021, or brain region, *F*(1, 96) = 0.15, *p* = .699, η²p = .002. The interaction between stimulation condition and brain region was also not significant, *F*(1, 96) = 0.10, *p* = .753, η²p = .001.

Change in positive mood on the VAMS showed a similar pattern. Mean positive mood change was −4.12 (SD = 56.74) in the PFC anodal condition, −15.64 (SD = 62.12) in the PFC sham condition, 1.36 (SD = 29.34) in the TPJ anodal condition, and −12.36 (SD = 37.92) in the TPJ sham condition. There was no significant main effect of stimulation condition, *F*(1, 96) = 1.70, *p* = .196, η²p = .017, or brain region, *F*(1, 96) = 0.20, *p* = .652, η²p = .002. There was also no significant stimulation condition × brain region interaction, *F*(1, 96) = 0.01, *p* = .910, η²p < .001. Overall, changes in VAMS negative and positive mood did not significantly differ between the experimental conditions.

### Adverse Effects

Adverse effects were also comparable across the experimental conditions. Mean adverse-effect scores were 4.48 (SD = 3.54) for PFC anodal stimulation, 4.04 (SD = 2.72) for PFC sham stimulation, 4.32 (SD = 3.35) for TPJ anodal stimulation, and 3.96 (SD = 2.56) for TPJ sham stimulation. A 2 × 2 ANOVA showed no significant main effect of brain region, *F*(1, 96) = 0.04, *p* = .845, η²p < .001, or stimulation condition, *F*(1, 96) = 0.42, *p* = .516, η²p = .004. The brain region × stimulation condition interaction was also not significant, *F*(1, 96) < 0.01, *p* = .948, η²p < .001, indicating that reported adverse effects did not significantly differ across conditions.

## Discussion

The present study examined whether focal stimulation of the left TPJ and dlPFC produced dissociable effects on source and reality monitoring. The findings provided partial support for the hypothesised target-specific effects of stimulation. Consistent with our prediction that TPJ stimulation would have a greater influence on reality monitoring, anodal left-TPJ stimulation reduced reality-monitoring sensitivity, whereas dlPFC stimulation had no significant effect on this measure. The prediction that dlPFC stimulation would facilitate source monitoring was also partially supported: anodal dlPFC stimulation was associated with a larger spoken–imagined difference in self–experimenter discrimination relative to sham stimulation. Although neither of the individual spoken or imagined comparisons was statistically significant, the significant context by stimulation interaction indicates that the effect of dlPFC stimulation depended on encoding context. In contrast, no equivalent context-dependent effect was observed following TPJ stimulation. Beyond these stimulation effects, source-monitoring performance was substantially better for spoken than imagined information, while reality-monitoring sensitivity was greater for self- than experimenter-generated negative information, providing additional evidence that the features of an episodic representation shape subsequent source and reality judgements.

The regional pattern of stimulation effects broadly supports, while also refining, the framework proposed by Perret et al. (2024). Their systematic review of non-invasive brain-stimulation studies identified lateral prefrontal and temporoparietal involvement in internal source monitoring and medial prefrontal and temporoparietal involvement in reality monitoring. The present findings provide clear convergence regarding the TPJ: anodal stimulation of the left TPJ reduced reality-monitoring sensitivity, consistent with Mondino et al. (2016), who similarly found that anodal left-TPJ stimulation impaired verbal reality monitoring and increased externalisation errors. Together, these findings suggest that increasing excitability within the left TPJ and its associated network can disrupt the discrimination of internally generated from externally perceived information (Mondino et al., 2016; Perret et al., 2024). In contrast, anodal dlPFC stimulation did not produce an overall change in reality-monitoring sensitivity, consistent with Perret et al.’s (2024) stronger association between reality monitoring and medial rather than lateral prefrontal regions. Thus, the present pattern supports the proposal that temporoparietal mechanisms contribute particularly to reality monitoring, while providing little evidence that the left dlPFC makes an equivalent contribution.

The source-monitoring findings are also broadly consistent with Perret et al. (2024), although they extend this framework in an important respect. Perret et al. (2024) classified internal source monitoring primarily in terms of discriminating between different internally generated sources, whereas the present task indexed agency by requiring participants to discriminate self- from experimenter-generated information across spoken and imagined contexts. Within this paradigm, the spoken–imagined difference in agent-discrimination sensitivity was larger following anodal than sham dlPFC stimulation, whereas no corresponding context by stimulation interaction was observed at the TPJ. This pattern was descriptively driven by greater self–experimenter discrimination for spoken items following anodal dlPFC stimulation, with no equivalent difference for imagined items, although neither individual simple effect was statistically significant following correction. Rather than indicating a general enhancement of source monitoring following dlPFC stimulation, these findings suggest that lateral prefrontal involvement may depend on the diagnostic features available within the retrieved episodic representation.

This interpretation is consistent with neuroimaging evidence implicating the left PFC in controlled source retrieval. Dobbins et al. (2002), for example, identified posterior dorsolateral and frontopolar activity associated with recollective monitoring during source-memory judgements. Spoken events in the present task contained multiple features that could inform agency judgements, including auditory and voice-identity information and, for self-generated speech, articulatory, motor, and proprioceptive information (Garrison et al., 2017; Palmeri et al., 1993). Imagined self and experimenter speech, by contrast, were both internally constructed and likely shared more similar imagery-related and cognitive features. Anodal dlPFC stimulation may therefore have influenced the strategic evaluation of diagnostic source features rather than source memory globally. This interpretation is consistent with the Source Monitoring Framework, according to which source attribution is inferred from the qualitative characteristics of retrieved memories rather than from explicit source labels (Johnson et al., 1993). Nevertheless, the absence of significant individual stimulation effects within the spoken and imagined conditions means that this mechanistic account should remain tentative.

The large behavioural effect of context further supports this account. Across stimulation and region conditions, agent-discrimination sensitivity was substantially greater for spoken than imagined information. Participants were therefore consistently better able to discriminate self- from experimenter-generated information when words had been spoken aloud. This advantage is consistent with spoken events providing richer and more distinctive information about their source than imagined events, in which representations of self and experimenter speech may share more similar phenomenological characteristics. By contrast, imagined self and experimenter speech were both internally generated and may therefore have shared more similar phenomenological characteristics. The magnitude of this effect demonstrates that source-monitoring performance depends strongly on the diagnostic information available within an episodic representation and provides a behavioural context for the context-dependent effect of dlPFC stimulation.

Emotional valence additionally influenced both mnemonic discrimination and response criterion. An agent by valence interaction was observed for reality-monitoring sensitivity, such that participants discriminated spoken from imagined events more accurately for self-generated negative words than experimenter-generated negative words, whereas no self– experimenter difference emerged for positive information. Self-generated negative words were also associated with a more liberal criterion for responding “spoken”. As the effect was evident in both sensitivity and criterion, however, the enhanced reality-monitoring performance for self-generated negative information cannot be explained solely by an increased tendency to classify these items as spoken. This pattern is consistent with evidence that negative and arousing information can enhance the encoding or retention of perceptual and contextual features that subsequently support discrimination between perceived and imagined events (Kensinger et al., 2007; Kensinger & Schacter, 2006).

The specificity of this effect to self-generated information may additionally reflect the social and embodied characteristics of the encoding episode. Saying a negative word aloud in the presence of the experimenter may have been more salient or arousing than either imagining oneself saying the word or hearing it spoken by another person. Although stimulus sets were matched on normative arousal, such ratings cannot capture context-dependent arousal elicited by producing emotionally salient language in an interpersonal setting. This additional salience may have strengthened auditory, articulatory, motor, and affective features of self-produced spoken events, increasing their discriminability from imagined events. This interpretation remains speculative, however, because subjective arousal during individual encoding conditions was not measured directly.

A related but distinct criterion effect emerged for source monitoring. For negative words, participants adopted a more liberal criterion for responding “self” following spoken than imagined encoding, whereas this pattern reversed for positive words. In the absence of a corresponding context by valence interaction in discrimination sensitivity, this effect suggests that context and valence altered the weighting of self- and other-response categories without changing participants’ ability to discriminate between agents. Such affective modulation of source-attribution criterion may be relevant to psychosis, where source-monitoring disturbances are frequently characterised by external attribution of self-generated events. For example, emotional salience has been associated with increased self-to-other misattributions in hallucination-prone nonclinical participants (Larøi et al., 2005) and external misattribution of negative self-generated words in psychosis (Costafreda et al., 2008). The direction of the present effect differed, with healthy participants more likely to attribute spoken negative information to themselves. One possibility is that socially salient self-production normally strengthens the weighting of self-agency cues, whereas this weighting is altered with psychosis vulnerability. Examining discrimination sensitivity and criterion separately across the psychosis continuum could help establish whether affect-related externalisation reflects degraded mnemonic discrimination, altered source-attribution criteria, or both.

Several limitations qualify the interpretation of the stimulation findings. Although the omnibus interactions demonstrated that stimulation effects differed according to cortical target and, for source monitoring, encoding context, the relevant simple effects were comparatively modest and some did not survive correction for multiple comparisons. The results should therefore not be interpreted as demonstrating a strict double dissociation between dlPFC and TPJ function. Moreover, anodal tDCS cannot be assumed to produce a simple increase in cortical activity and behavioural effects may reflect changes within distributed networks connected to the stimulated target rather than modulation of the target region in isolation.

Nevertheless, the use of a focal montage, sham control, independent manipulation of two cortical targets, and current-flow modelling provides greater anatomical specificity than conventional bipolar stimulation designs. Future work should replicate these effects using converging methods and directly examine whether stimulation alters the retrieval or weighting of specific perceptual, motor, and affective features of episodic memories.

In conclusion, the present findings provide partial causal support for dissociable contributions of lateral prefrontal and temporoparietal systems to source and reality monitoring. The detrimental effect of left-TPJ stimulation on reality-monitoring sensitivity broadly converges with previous stimulation evidence and the neural framework synthesised by Perret et al. (2024), whereas the context-dependent effect of dlPFC stimulation suggests that lateral prefrontal contributions to source attribution may depend on the contextual features available within a remembered event. More broadly, the findings suggest that source and reality monitoring depend on how neural systems retrieve and evaluate the perceptual, agent-related, and affective characteristics that specify the origin of an episodic memory.

## Acknowledgements

We thank all participants for their time and effort

## Data availability

Data will be available upon request

