## Supplementary for "Causal contributions of the dorsolateral prefrontal cortex and temporoparietal junction to source and reality monitoring"

**Supplementary Table 1**

*Mixed ANOVA results for source-monitoring discrimination sensitivity (d′)*

| **Effect** | **F(1, 96)** | **p** | **η²G** |
| --- | --- | --- | --- |
| Region | 0.32 | .574 | .002 |
| Stimulation | 0.11 | .743 | < .001 |
| **Region × Stimulation** | **4.39** | **.039** | **.023** |
| **Context** | **205.80** | **< .001** | **.280** |
| Context × Region | 0.46 | .500 | < .001 |
| Context × Stimulation | 1.29 | .258 | .002 |
| **Context × Region × Stimulation** | **6.03** | **.016** | **.011** |
| Valence | < 0.01 | .968 | < .001 |
| Valence × Region | 0.17 | .680 | < .001 |
| Valence × Stimulation | 0.07 | .789 | < .001 |
| Valence × Region × Stimulation | 0.90 | .346 | .002 |
| Context × Valence | 2.23 | .138 | .003 |
| Context × Valence × Region | 2.02 | .159 | .003 |
| Context × Valence × Stimulation | 0.12 | .731 | < .001 |
| Context × Valence × Region × Stimulation | 0.73 | .394 | < .001 |

*Note.* η²G = generalised eta-squared. Statistically significant effects (p < .05) are shown in bold.

**Supplementary Table 2**

*Mixed ANOVA results for source-monitoring response criterion (c)*

| **Effect** | **F(1, 96)** | **p** | **η²G** |
| --- | --- | --- | --- |
| Region | 0.17 | .685 | < .001 |
| Stimulation | 0.07 | .795 | < .001 |
| Region × Stimulation | 0.12 | .734 | < .001 |
| Context | 0.79 | .377 | .002 |
| Context × Region | 0.01 | .912 | < .001 |
| Context × Stimulation | 2.09 | .151 | .005 |
| Context × Region × Stimulation | 0.15 | .695 | < .001 |
| **Valence** | **3.96** | **.049** | **.008** |
| Valence × Region | 1.29 | .259 | .003 |
| Valence × Stimulation | 1.93 | .168 | .004 |
| Valence × Region × Stimulation | 0.01 | .905 | < .001 |
| **Context × Valence** | **28.10** | **< .001** | **.047** |
| Context × Valence × Region | 3.47 | .066 | .006 |
| Context × Valence × Stimulation | 1.07 | .304 | .002 |
| Context × Valence × Region × Stimulation | 0.26 | .611 | < .001 |

*Note.* η²G = generalised eta-squared. Statistically significant effects (p < .05) are shown in bold.

**Supplementary Table 3**

*Mixed ANOVA results for reality-monitoring discrimination sensitivity (d′)*

| **Effect** | **F(1, 96)** | **p** | **η²G** |
| --- | --- | --- | --- |
| Region | 0.71 | .402 | .004 |
| Stimulation | 0.80 | .374 | .005 |
| **Region × Stimulation** | **4.20** | **.043** | **.025** |
| **Agent** | **41.61** | **< .001** | **.065** |
| Agent × Region | 0.57 | .454 | < .001 |
| Agent × Stimulation | 1.51 | .223 | .002 |
| Agent × Region × Stimulation | 0.05 | .829 | < .001 |
| Valence | 0.80 | .372 | .001 |
| Valence × Region | 0.13 | .716 | < .001 |
| Valence × Stimulation | 2.08 | .152 | .003 |
| Valence × Region × Stimulation | 0.10 | .751 | < .001 |
| **Agent × Valence** | **29.11** | **< .001** | **.035** |
| Agent × Valence × Region | 0.41 | .525 | < .001 |
| Agent × Valence × Stimulation | 0.03 | .863 | < .001 |
| Agent × Valence × Region × Stimulation | 1.28 | .261 | .002 |

*Note.* η²G = generalised eta-squared. Statistically significant effects (p < .05) are shown in bold.

**Supplementary Table 4**

*Mixed ANOVA results for reality-monitoring response criterion (c)*

| **Effect** | **F(1, 96)** | **p** | **η²G** |
| --- | --- | --- | --- |
| Region | 0.10 | .756 | < .001 |
| Stimulation | 1.24 | .269 | .007 |
| Region × Stimulation | 3.09 | .082 | .017 |
| Agent | 1.22 | .272 | .002 |
| Agent × Region | 0.03 | .863 | < .001 |
| Agent × Stimulation | 0.36 | .550 | < .001 |
| Agent × Region × Stimulation | 1.64 | .203 | .003 |
| Valence | 3.40 | .068 | .006 |
| Valence × Region | 0.15 | .697 | < .001 |
| Valence × Stimulation | 0.76 | .384 | .001 |
| Valence × Region × Stimulation | 0.04 | .846 | < .001 |
| **Agent × Valence** | **14.87** | **< .001** | **.018** |
| Agent × Valence × Region | 0.99 | .322 | .001 |
| Agent × Valence × Stimulation | 0.91 | .343 | .001 |
| Agent × Valence × Region × Stimulation | 0.28 | .598 | < .001 |

*Note.* η²G = generalised eta-squared. Statistically significant effects (p < .05) are shown in bold.
